# A Data-Driven Closed-Loop Control Approach to Drive Neural State Transitions for Mechanistic Insight

**DOI:** 10.1101/2025.07.21.665992

**Authors:** Niklas Emonds, Evelyn Herberg, Martin Fungisai Gerchen, Marc Pritsch, Joshua Rocha, Vera Zamoscik, Peter Kirsch, Roland Herzog, Georgia Koppe

## Abstract

Repetitive negative thinking (RNT) is a transdiagnostic risk factor for mood disorders, consistently associated with altered biological substrates, including functional connectivity in key brain networks. As a stable cognitive feature linked to vulnerability across disorders, RNT presents a compelling target for intervention. However, leveraging RNT as a modifiable mechanism requires a deeper understanding of its causal neural dynamics and how targeted modulation can induce adaptive change.

We introduce a data-driven framework that combines dynamical system reconstruction (DSR) with model predictive control (MPC) to infer optimal control policies for transitions between resting and sad mood brain states from functional magnetic resonance imaging (fMRI) data. Using nonlinear generative DSR models trained on individuals with remitted major depressive disorder (rMDD) and matched healthy controls (HCs), we derive region-specific, state-dependent control strategies. We find that small brain regions (e.g., sgACC, NAcc) exhibit higher controllability, requiring less energy to drive state transitions.

Critically, rMDD participants require less control energy than HCs to move *into* sad mood from rest and - unexpectedly - also to move *back* to rest, though the latter effect is spatially restricted. Despite comparable target attainment, rMDD participants remain closer to the sad mood distribution when returning to rest, indicating a residual negative-affect bias. Our data-driven analysis reveals elevated effective coupling in rMDD, most prominently toward (but not away from) the DLPFC. Across regions, greater coupling is associated with reduced control energy, suggesting that enhanced network influence facilitates more efficient state transitions.

These results suggest dynamics in rMDD that facilitate entry into negative affect and hinders full disengagement without sustained input, highlighting closed-loop control as a tool for mechanistic insight and potentially for designing targeted neuromodulatory interventions in the future.

## Introduction

Repetitive negative thinking (RNT)—a pattern of persistent, repetitive negative thoughts from which it is difficult to disengage—is a core transdiagnostic feature of mood and anxiety disorders (Bell et al., 2023; Moulds and McEvoy, 2025). It encompasses cognitive phenomena such as rumination and worry, and is characterized by its abstract, self-focused content and perceived uncontrollability (Gustavson et al., 2018). Importantly, RNT often persists beyond symptomatic remission and has been robustly linked to the onset, maintenance, and severity of psychopathology (Spinhoven et al., 2018; Timm et al., 2017). Intervening on RNT thus remains a pressing goal for prevention and treatment. However, effective intervention requires mechanistic insight into how neural systems shift into and out of RNT-associated states.

Previous studies have associated RNT with altered functional connectivity across large-scale brain networks, particularly increased coupling between default mode and fronto-parietal regions and decreased coupling with the salience network (Lydon-Staley et al., 2019; Zamoscik et al., 2018; Zhu et al., 2012). Elevated levels of connectivity involving the parahippocampal gyrus (PHG), anterior cingulate cortex (ACC), and posterior cingulate cortex (PCC) have been especially prominent in remitted depression (Zamoscik et al., 2014). Yet these descriptive findings fall short of revealing the causal mechanisms that govern transitions between healthy and maladaptive cognitive states.

To uncover such mechanisms, generative models that simulate brain dynamics—and allow for controlled perturbation—have emerged as a powerful framework (Durstewitz et al., 2021, 2023; Luo et al., 2025). Within this framework, optimal control theory (Brunton and Kutz, 2019) enables formal inference of the minimal interventions required to shift brain states under physiological constraints. Prior work has applied linear control models to empirical connectivity data to study cognitive transitions, predict symptom severity, or evaluate controllability in neurological conditions (Betzel et al., 2016; Deng et al., 2022; Lynn and Bassett, 2019; Medaglia et al., 2017; Parkes et al., 2024; Yao et al., 2025; Zhou et al., 2025). Others have introduced biophysical or neural field models to simulate state transitions in conditions such as epilepsy (Costa et al., 2024; Deco et al., 2019; Salfenmoser and Obermayer, 2022; Taylor et al., 2015). In mental disorder treatments, interventions have also been conceptualized as model predictive control problems (Brar et al., 2018; Chang et al., 2020; Santaniello et al., 2011; Steffen et al., 2024). While valuable, many of these approaches rely on simplified, often linear dynamics, which fail to capture phenomena such as multistability or chaos that characterize real neural systems (Breakspear, 2017; Driscoll et al., 2024; Strogatz, 2015; Volkmann et al., 2024), and therefore limit mechanistic interpretations. Models with greater biological detail often rely on handcrafted equations (Martínez et al., 2023; Taylor et al., 2015), which are typically not learned from data and are thus difficult to adapt or generalize across individuals and datasets.

Recent advances in dynamical systems reconstruction (DSR) offer a promising alternative. Data-driven methods learn expressive, nonlinear models directly from time series data and can act as computational surrogates of neural systems (Brenner et al., 2025; Brunton and Kutz, 2019; Durstewitz et al., 2023; Hess et al., 2023; Pathak et al., 2018). Here, we build on these advances by combining nonlinear DSR with optimal control to investigate the mechanisms underlying RNT. Using fMRI data from participants with a remitted major depressive disorder (rMDD) and matched healthy controls (HC) during a sad mood induction task, a paradigm eliciting RNT, we train DSR models that capture individual neural dynamics (Brenner et al., 2025; Volkmann et al., 2024). By deriving and analyzing optimal control policies within these models, we aim to identify the causal structure that enables transitions between healthy and pathological states.

Our contributions are fourfold: We present a novel data-driven framework for inferring closed-loop optimal control policies based on DSR models. We perform in silico experiments to examine control trajectories between RNT-associated and baseline brain states, targeting single brain regions. We show that inferred control policies differ between rMDD and HC individuals, reflecting altered neural dynamics. We identify key brain regions with elevated controllability and link these findings to underlying connectivity. Together, these results offer a mechanistic perspective on RNT and establish a foundation for future causal interventions targeting pathological brain states.

## 2 Results

This study seeks to derive optimal control policies from fMRI recordings collected from rMDD and HC individuals to gain mechanistic insights into the neural processes underlying transitions between resting and sad mood induction states. Figure 1 depicts the general procedure.

**Figure 1:**
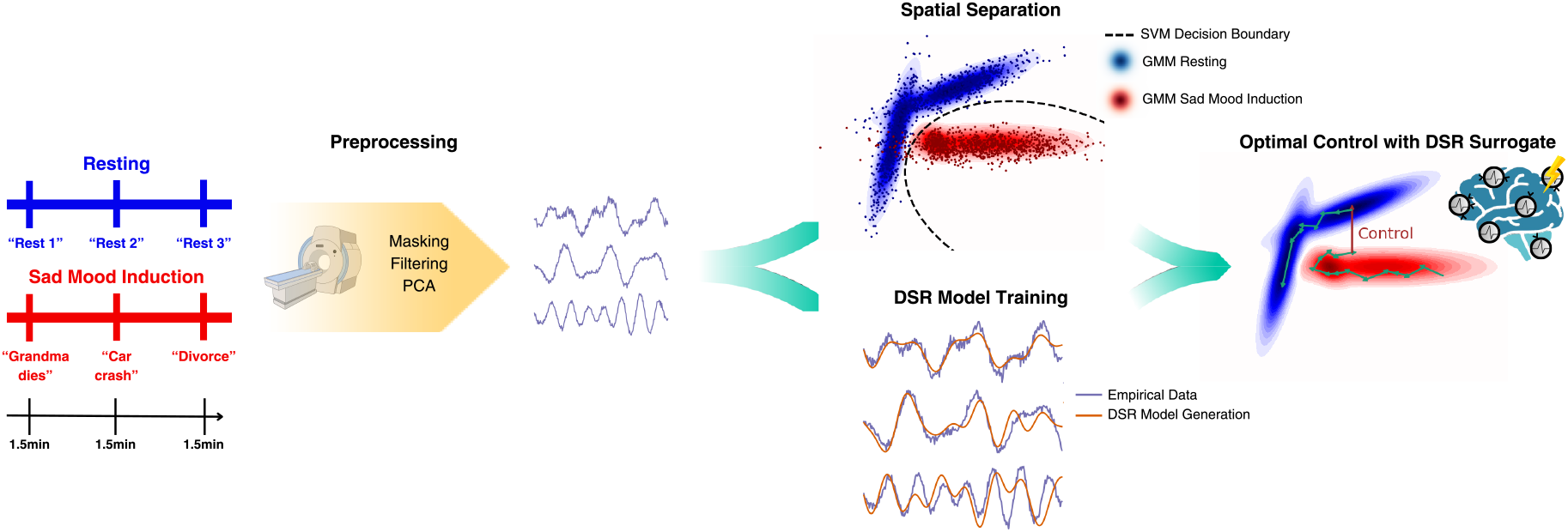
Neural activity was measured during two cognitive states, a resting state, and a sad mood induction state (Zamoscik et al., 2014). After preprocessing, fMRI recordings are used to train a classifier that separates resting from sad mood (top), and to train DSR models that serve as brain surrogates (bottom). The trained classifiers serve to identify the control target. After verifying the reconstruction quality, we train optimal control strategies on the inferred DSR models that drive neural states from resting to sad mood induction states.

### Spatial Separation Between Resting and Sad Mood Induction

To characterize differences in the representation and distribution of the two cognitive states—needed for downstream control analyses—we first trained separate Gaussian Mixture Models (GMMs) for the resting state and sad mood induction conditions (see Figure 2a and Section 5.2.1 for details on the preselected regions of interest; ROIs). Consistent with both groups reporting increased negative affect after sad mood induction (Zamoscik et al., 2014), the two states were well separable, with the GMMs achieving high classification accuracy on 25% held-out test sets using 5-fold cross-validation (94%±3%), by assigning each test sample to the condition with higher likelihood (Figure 2b). Comparable classification performance was achieved using support vector machines (SVMs) with a quadratic polynomial kernel (93%±3%); in contrast, linear classifiers such as linear discriminant analysis (LDA) and linear-kernel SVMs performed poorly, highlighting the nonlinear nature of the boundary separating the two cognitive states. No significant group differences in classification performance were observed between HC and rMDD participants. GMMs were used in downstream analyses to define targets for optimal control and to decode neural activity, inferring cognitive states from both empirical and generated trajectories. Neural states remained separable when sampling from the GMMs (Figure A2a).

**Figure 2:**
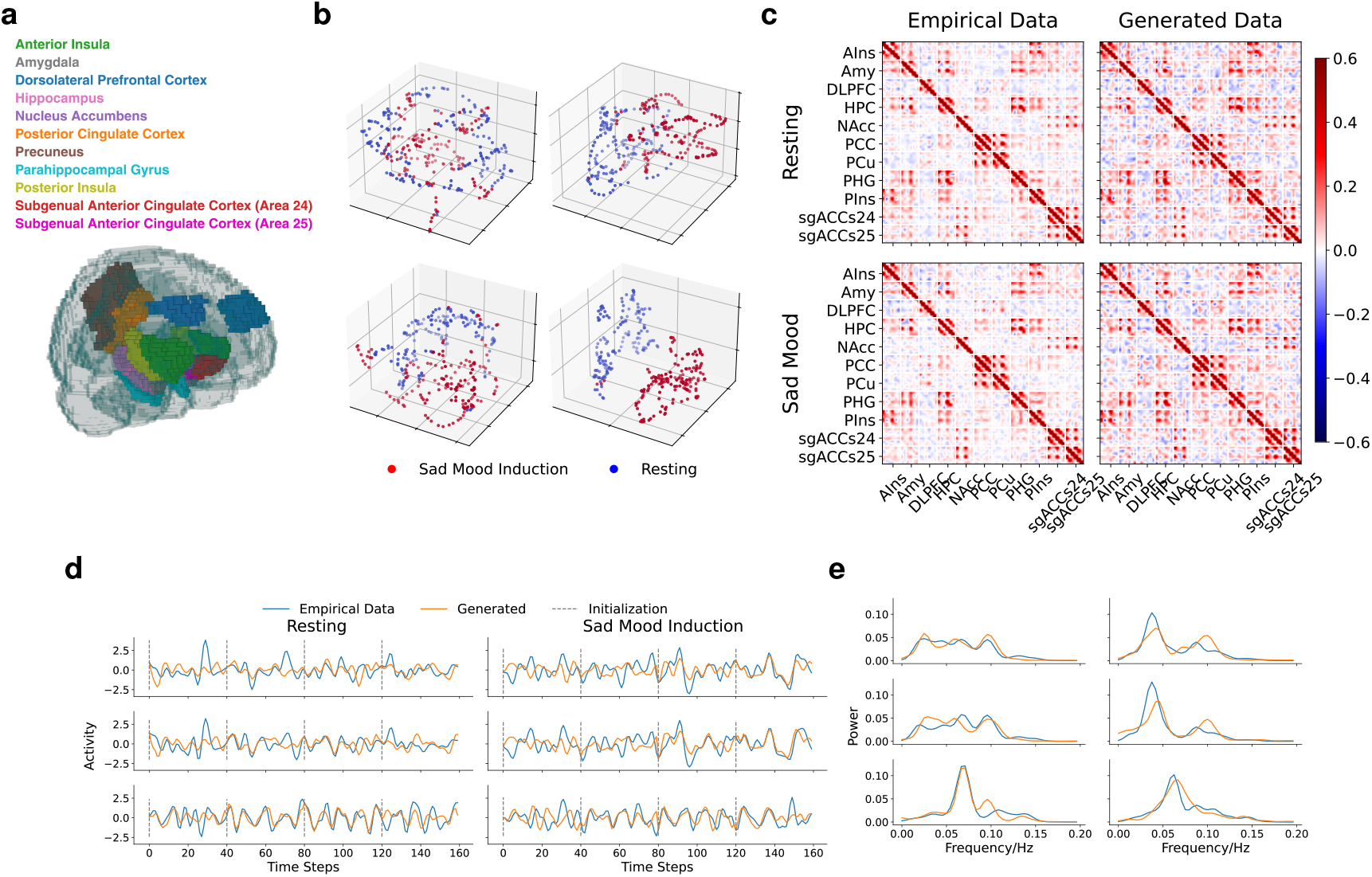
Neural State Classifier and DSR Model Performance. **a**: Eleven bilateral regions of interest (ROIs) implicated in depression, as visualized here, were selected for analysis. **b**: Neural state representations projected via UMAP (McInnes et al., 2018), color-coded by condition (resting: blue; sad mood induction: red) for selected participants. **c**: Median cross-correlation matrices between selected ROIs for empirical test data (left), and simulated test data from the DSR models (right). The observed connectivity patterns are consistent across empirical and simulated data. **d**: Example time series of empirical neural activity (blue) and model reconstructions (orange) for a few representative time series. Grey dashed lines indicate the initialization period for the model-generated trajectories. The final 40 time steps (test set) were not used during training. **e**: Normalized power spectra of the time series shown in **d**.

### DSR Models Learn the Underlying Dynamics

Next, we inferred subject-level generative models from the neural data using piecewise linear recurrent neural networks (PLRNNs) trained with a state-of-the-art DSR protocol (Hess et al., 2023). These state-space type models—also referred to as *world models* in model-based reinforcement learning (RL; Ha and Schmidhuber, 2018; Hafner et al., 2025)—consist of a latent PLRNN model coupled to a linear decoder, enabling the reconstruction of high-dimensional neural trajectory dynamics (see Section 5.3 for details). Representative reconstructions from the inferred models are shown in Figure 2d.

The inferred dynamics were found to be chaotic, characterized by positive maximum Lyapunov exponents (*λ*_max_ = 0.1±0.02), in line with previous findings on neural recordings during resting states (Volkmann et al., 2024). This chaotic nature makes it inherently difficult to achieve a point-by-point match between predicted and empirical trajectories over time (see also Koppe et al., 2019; Mikhaeil et al., 2022; Schmidt et al., 2021). Despite this, the models successfully capture the data’s underlying temporal structure, as evidenced by strong alignment in the recovered frequencies, measured in terms of the Hellinger distance *D*_*H*_ between power spectra (*D*_*H*_ = 0.22 ± 0.04, Figure 2e). Moreover, the cross-correlation between different brain regions—commonly referred to as functional connectivity—exhibits highly similar patterns in both the empirical data and the model-generated test set trajectories (Figure 2c). These results underscore that, although exact trajectory matching may not be expected, the inferred models nonetheless reproduce the essential statistical and spectral properties of the observed neural activity, indicating successful system identification under the DSR framework (Durstewitz et al., 2023).

### Learning Successful Control Policies

Having established our DSR models, we next inferred optimal control policies by augmenting the models with an additive nonlinear control term (see Section 5.3). To derive these policies, we fixed the DSR model parameters and optimized only the control input. The objective of this optimization was to steer the inferred systems toward a desired target in a state-dependent manner. In other words, the resulting policies specify how specific brain regions should be stimulated depending on the current state, in order to reach a given target (e.g., how the amygdala must be modulated in order for the dynamical system to transition from the resting to sad mood state—or vice versa—in a given time). Notably, our framework allows control to be restricted to specific output dimensions, such as, in this case, individual brain regions.

We considered a *soft target* strategy, which assigns probabilities across states based on the above mentioned GMMs and aims to steer the system toward probabilistically representative states (see Section 5.3 for details). To assess the effectiveness of the control policies, we evaluated the negative log-likelihood (NLL_Ind_ and NLL_Rest_ for the sad mood induction and resting state, respectively) under the GMM. The inferred control policies are governed by two hyperparameters: the control energy regularization parameter *λ*_E_, which penalizes the magnitude of the control input, and the temporal horizon *D*, which defines the number of time steps allowed for the system to reach the target (cf. eqn. (4)).

To determine appropriate parameter values for the control energy regularization term *λ*_E_, we first investigated the effect of *λ*_E_ on the trade-off between control energy and proximity to the target, for models trained using the soft target approach on the bilateral amygdala (Amy) (Figure 3a) and transitions from resting to sad mood. The resulting curve shows that for sufficiently large (or small) values of *λ*_E_, the optimization favors minimizing either energy or target proximity, respectively. Importantly, the curve exhibits an elbow around *λ*_E_ = 0.1, beyond which further increases in control energy do not improve target proximity—indicating a balance point in the trade-off. Based on this observation, we fixed *λ*_E_ at 0.1 for all subsequent analyses. The temporal horizon *D* was set to 10 to give the network some time to move toward the target using its own dynamics, while still requiring it to reach the target within a reasonable time frame for fMRI. We report results for alternative *λ*_E_ and *D* in the Supplement (Figure A3), demonstrating that our main conclusions are insensitive to moderate hyperparameter changes.

**Figure 3:**
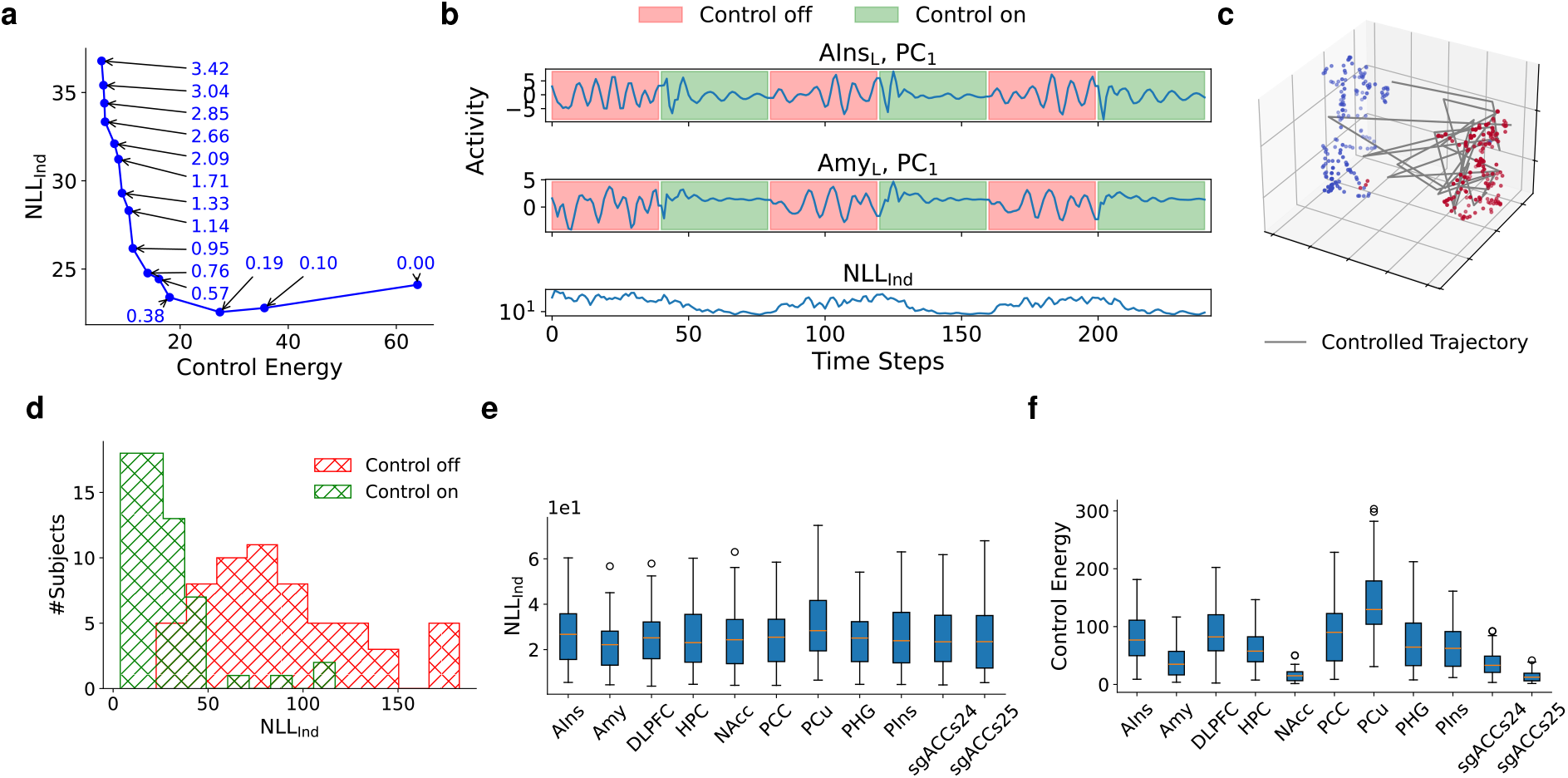
Control Succesfully Steers the Dynamics. **a**: Pareto front illustrating the trade-off between NLL_Ind_ and control energy in a multiobjective optimization, using the amygdala as an example. Corresponding values of *λ*_E_ are indicated in the diagram. Saturation of the control occurs at approximately *λ*_E_ = 10^™1^, beyond which less regularization no longer improves NLL_Ind_. **b**: Example trajectories from a controlled region (left Amy; middle) and an uncontrolled region (left anterior insula; AIns; top), both move toward the sad mood induction state, as confirmed by decoding with a Gaussian Mixture Model (bottom). **c**: UMAP projection reveals that generated trajectories align closely with the empirical sad mood induction state distribution (red), while diverging from resting state distributions (blue). **d**: Across all 60 subjects, control significantly reduces the NLL of the trajectory over time steps under the GMM. **e**: Steering accuracy, quantified as the NLL_Ind_ target proximity (y-axis), is comparable across all stimulation sites (x-axis), indicating that each region can drive the system equally close to the desired state. **f** : In contrast, the control energy (y-axis) required to achieve that steering differs significantly between regions (x-axis).

Figure 3b-3e illustrates the efficacy of our control protocol for the case of control towards sad mood induction states, and Figure A4 visualizes control policies across regions for a selected individual. Upon control activation, neural activity converges to the target, reverting to its intrinsic chaotic dynamics once control is deactivated. While state separation became more pronounced when control was applied across multiple brain regions (Figure A5), we observed a significant reduction in target proximity even when controlling single regions. This improvement is driven by the indirect modulation of non-controlled regions through network-level interactions. The control policies also demonstrated the ability to generalize to previously unseen states: although the system may encounter novel configurations during deployment, the policies consistently guided the dynamics back toward the target. In principle the same procedure can be applied using a hard target approach (Figure A5).

### Small Brain Regions Exhibit High Controllability

For subsequent analyses, we focused on two key questions: (1) Can we identify brain regions that are more effective than others in driving state transitions? and (2) How do these effects differ between individuals with rMDD and HC? Using the soft target approach, we derived separate control policies for each bilateral ROI.

While control policies across all regions achieved a comparable proximity to the target (Figure 3e), the amount of energy required to achieve this convergence varied between regions (Figure 3f). Smaller regions, such as the subgenual ACC (sgACC) and nucleus accumbens (NAcc), exhibited enhanced controllability, reaching the target more efficiently. Less controllability in larger brain areas might arise because they exhibit more spatial variability (correlation between size and total variance *r*(11) = 0.97, *p <* 0.001), causing the first principal components (PCs) to have greater variance as confirmed by a strong correlation between brain region size and control energy (*r*(11) = 0.93, *p <* 0.001). This increased dynamic range may require greater control energy to steer the distributed activity toward the target. On the other hand, sgACC and NAcc play a central role in negative mood control and are established targets for deep brain stimulation (Figee et al., 2022), such that increased controllability may also be attributed to an emphasized functional role.

### Asymmetric Neural Controllability in rMDD: Reduced Energy Costs and Residual Bias Toward Sad Mood States

Next, we examined differences in control policies between rMDD and HCs during the transition from resting to sad mood induction states and vice versa.

During transitions *from the resting state to sad mood* (Figure 4a), individuals with rMDD required significantly less control energy than HCs across nearly all examined regions, with the exception of the NAcc and PCC (Figure 4b). This reduction in energetic cost indicates enhanced neural controllability in rMDD, suggesting that both the transition to and maintenance of the sad mood state were more easily achieved in this population. Importantly, this effect was robust across a broad range of regularization parameters and could not be explained by differences in the separability of baseline proximity between the resting and sad mood states (Figure A2c). Despite the energetic differences, no significant group differences were observed in the proximity to the (sad mood) target state as measured by NLL_Ind_ (Figure 4c), nor to the initial (resting) state (Figure A6a).

**Figure 4:**
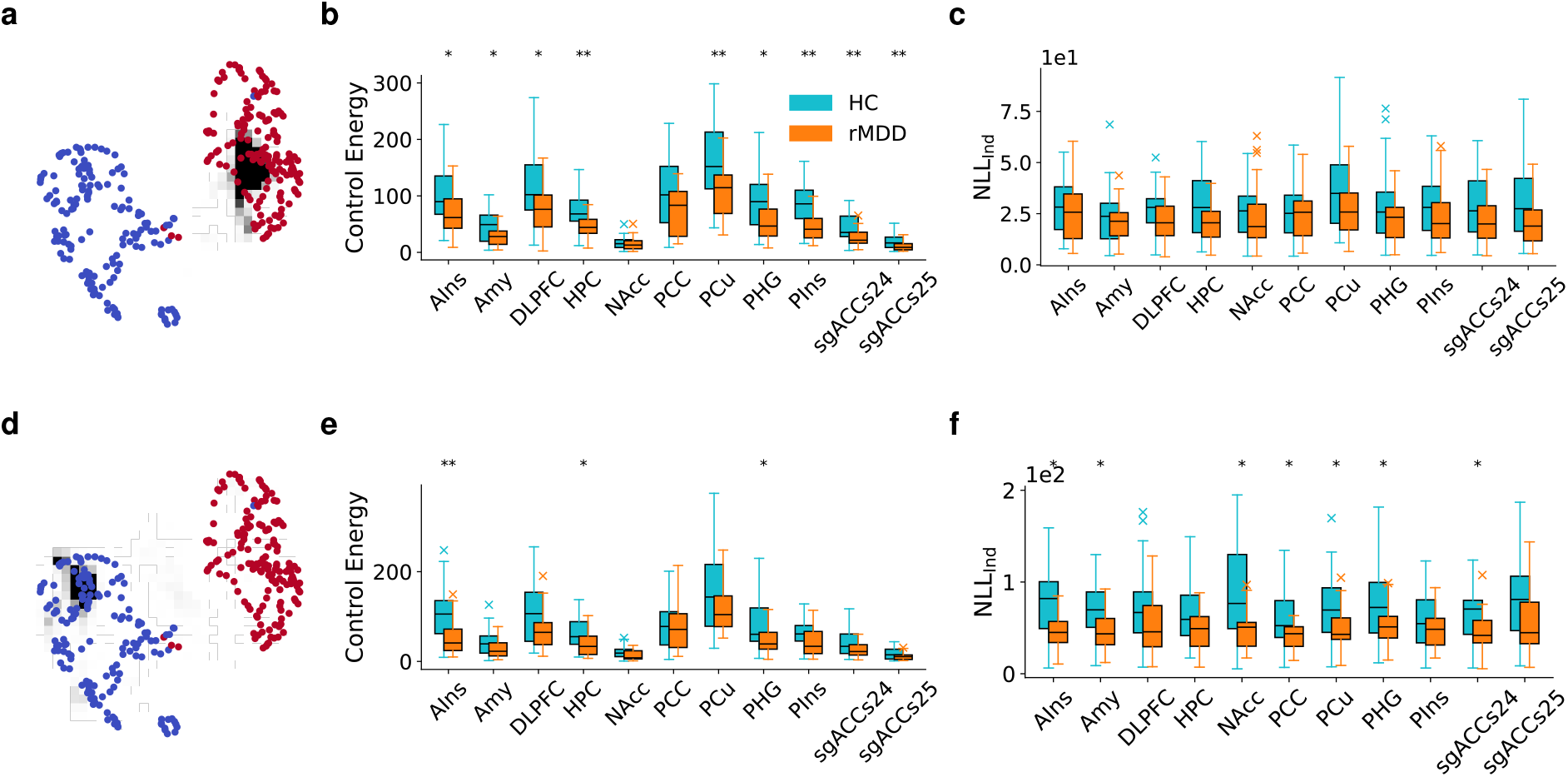
Group Control Analyses. **a**: UMAP visualization of empirical states (blue and red). Different initializations of controlled trajectories yield the heat map (black) of the controlled activity which shows that the activity remains in the sad mood induction region due to the control. **b**: Control energy required to drive the brain toward the sad mood state by stimulating individual brain regions. rMDD participants consistently require less energy than HC. **c**: Despite the different amounts of energy expended, the groups do not show significant differences in proximity to the sad mood states. **d, e, f** : Same as in **a, b, c**, but for control towards resting state. *p* values reflect FDR-corrected significance thresholds at thresholds ^*^*p <* 0.05, ^**^*p <* 0.01.

When transitioning *from the sad mood state back to resting state* (Figure 4d), rMDD participants again showed a tendency toward reduced control energy, though this effect was more spatially restricted, reaching statistical significance only in the AIns, HPC, and PHG (Figure 4e and A8). While both groups approached the target with comparable proximity (Figure A6b), rMDD individuals remained closer to the sad mood state (Figure 4f), suggesting a lingering bias toward the negative-affect configuration despite reduced energetic demands. This was found to be significant for most regions, apart from the dorsolateral prefrontal cortex; (DLPFC), hippocampus (HPC), posterior inusla (PIns) and parts of sgACC).

### Connectivity Predicts Controllability

Finally, we sought to identify mechanistic differences between groups that may underlie the altered controllability patterns observed in rMDD. To this end, we leveraged the piecewise linear architecture of our data-driven DSR model to estimate *coupling matrices C*, that is, transition dynamics expressed in observation space (see Section 5.3.5 for details).

The spectral norm of these coupling matrices, given by the largest singular value, was significantly higher in the rMDD group (*p <* 0.001, Figure 5a), indicating stronger global dynamical coupling relative to HCs.

**Figure 5:**
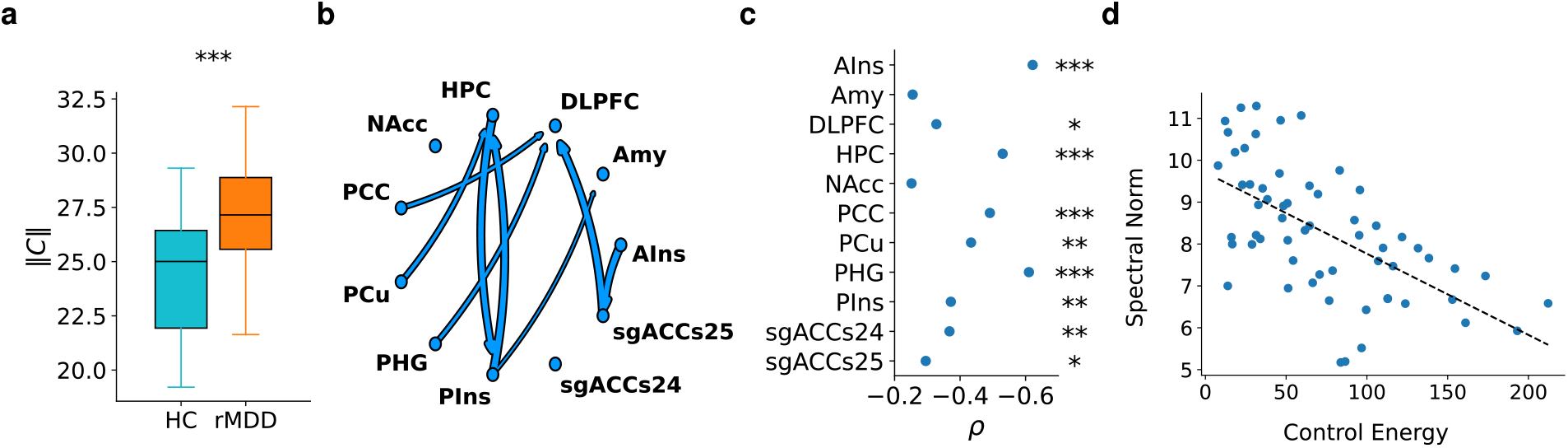
Brain Region Coupling. **a**: Global coupling strength, measured as the spectral norm of the full coupling matrix *C*, is significantly higher in rMDD. **b**: Pairwise coupling strength between brain regions is computed as the spectral norm of 6 × 6 submatrices. Significant differences between groups in directed coupling (*p*_FDR_ *<* 0.05), depicted as arrows indicating the direction and strength of influence from one region to another. **c**: Correlation between regional coupling strength and control energy across subjects. Significant negative correlations indicate that stronger outward influence is associated with less control effort to reach the sad mood states, suggesting a mechanistic link between connectivity and controllability in rMDD. **d**: PHG coupling strength (as measured via spectral norm) versus required control energy per subject.

To localize these effects, we examined the (6×6) (each region is associated with 2 hemispheres and 3 PCs) submatrices of the full coupling matrices that quantify how activity in one region influences transitions in another. This revealed elevated directed coupling in rMDD, particularly toward—but not away from—the DLPFC (Figure 5b), a region frequently implicated in cognitive control and the top-down regulation of affect.

Finally, we asked whether regional coupling strength predicts controllability. Specifically, we computed the spectral norm of the 66×6 submatrices representing directed coupling from each region to all others, and correlated these values with the control energy required to reach the sad mood target (Figure 5c, 5d). Across most regions, stronger outward coupling was associated with lower control energy, suggesting that regions with greater influence over the network facilitate more efficient control. This effect was particularly pronounced in AIns, HPC, PCC, and PHG — all of which showed increased coupling in rMDD, and many of which directly or indirectly modulate DLPFC dynamics (Figure 5b). The correlation was preserved for the reverse direction (Figure A7a and A7b). These findings may indicate that regions exerting strong outbound influence, especially those interfacing with prefrontal control, play a central role in steering brain state transitions in depression.

## 3 Discussion

This study introduces a data-driven framework that combines DSR with model predictive control (MPC) to examine how brain dynamics in remitted rMDD support or resist transitions between emotional states. Our approach leverages expressive generative models, trained solely on empirical fMRI time series, to reconstruct neural dynamics and enable in silico causal intervention analyses. By treating these models as computational surrogates of the brain, we extend prior control-theoretic work — often limited to linear (Deng et al., 2022; Gu et al., 2015; Parkes et al., 2024; Yao et al., 2025) or biophysically constrained (Martínez et al., 2023; Taylor et al., 2015) models — into a generalizable and expressive machine learning framework capable of simulating and controlling neural trajectories at the level of individual subjects.

Unlike traditional correlational analyses, our framework provides a causal account of how neural perturbations drive transitions between functional states. By learning optimal control policies on reconstructed latent dynamics, we can identify how activating or deactivating specific brain regions shapes trajectories of neural activity. We illustrate this approach by examining mechanisms through which individuals with rMDD more readily enter, sustain, or exit states of negative affect, highlighting potential intervention targets for modulating pathological state transitions.

### Validity of the Generative Models

Central to our approach is the training of DSR models that accurately reconstruct empirical neural dynamics. Beyond capturing the complex temporal evolution of the data, the models replicated key statistical and connectivity-based properties. In particular, functional connectivity patterns in the model-generated trajectories closely mirrored those observed in the empirical recordings. Notably, the models revealed elevated directed coupling in rMDD participants, consistent with previous findings of altered network interactions in depression (Lydon-Staley et al., 2019; Zhu et al., 2012), and elevated connectivity observed specifically in this dataset (Zamoscik et al., 2014), although our results were derived entirely from a data-driven approach. This cross-methodological convergence underscores the robustness of both our modeling framework and its capacity to extract clinically relevant neural signatures.

### Reduced Control Energy and Persistent Sad-Mood Dynamics in rMDD

Our first finding indicates that individuals with rMDD require significantly less control energy to transition into and maintain sad mood states compared to HCs. This reduced energetic cost suggests that, in rMDD, the dynamical landscape is shaped in a way that favors transitions toward negative affect, aligning with prior reports of heightened negative and diminished positive affect in response to sad mood induction in this group (Zamoscik et al., 2014). We also found that this energetic advantage was widespread across regions, with only a few exceptions. Such global efficiency implies that the facilitation is not localized to a single structure but emerges from network-level interactions.

More unexpectedly, rMDD participants also required less energy than controls to transition from the sad mood state back to rest (although this effect was far less pronounced across regions). This might appear inconsistent with the idea of heightened vulnerability; however, a closer look revealed an important asymmetry. Although both groups could be guided toward the resting-state distribution under active control, rMDD trajectories *remained closer to the sad-mood state* even while control was still applied. This implies that, in rMDD, even when the system is successfully pushed toward rest, its underlying dynamics continue to partially align with the sad-mood configuration. The models therefore indicate a form of incomplete disengagement: the system can move toward rest efficiently, but it does not fully leave the negative-affect manifold. This may reflect a form of neural hysteresis: the system can be directed back, but it resists complete reconfiguration and continues to favor the previously occupied mood state. Such persistence could provide a mechanistic explanation for the well-documented difficulty rMDD individuals have in sustaining positive or neutral affect even after recovery from depressive episodes (Moulds and McEvoy, 2025, Paykel, 2008; Timm et al., 2017).

### Connectivity as the Mechanistic Link

The reduced control energy in rMDD was strongly predicted by increased coupling strength across the network. Regions with stronger outbound influence over others required less external input to drive the whole system toward the target. This finding provides a mechanistic bridge between controllability and network structure: highly connected regions act as natural leverage points that can efficiently propagate state changes. It is also consistent with the observation that networks with higher mean degree not only need fewer driver nodes but also incur lower control energy, because the extra connections provide multiple, “low-resistance” pathways for steering the system (Liu et al., 2011). Interestingly, we observed elevated coupling especially directed toward (and not away from) the DLPFC—a region implicated in cognitive control and emotion regulation (e.g., Nejati et al., 2021; Vanderhasselt et al., 2013). This asymmetry may reflect a compensatory reorganization in which affective regions exert greater influence over prefrontal circuits, potentially undermining top-down control and contributing to the persistence of negative mood states observed in rMDD.

### Outlook

Our findings have at least two implications. First, they offer a mechanistic explanation for the increased vulnerability to negative affect in (remitted) depression: the brain’s intrinsic dynamics make transitions into sadness easy and recovery only partially complete, even under active control. Second, they suggest putative causal targets for intervention. Our analyses revealed strong negative correlations between control energy and coupling strength in multiple brain regions examined, but particularly the HPC, the PHG and the AIns—regions already implicated in affective control and neuromodulation therapies (Arulchelvan and Vanneste, 2023; Liu et al., 2021; Roet et al., 2020). Moreover, small regions like sgACC and NAcc were confirmed as particularly efficient hubs facilitating transitions between resting and sad mood states, emhphasizing their role as efficient control points for modulating maladaptive dynamics in depressive disorders.

Beyond theoretical insight, our framework offers immediate translational opportunities. Because control is learned separately for each individual brain region, the method provides a principled way to assess regional controllability—an insight that can be used for personalized intervention planning. In particular, the framework allows for in silico perturbation analyses of potential stimulation targets, identifying regions that can steer the system with minimal energy or maximal precision (see also Fechtelpeter et al., 2025; Luo et al., 2025; Muldoon et al., 2016). Coupled with non-invasive brain stimulation technologies—such as transcranial magnetic stimulation or transcranial direct current stimulation—our framework could serve as the computational core of closed-loop neurostimulation systems. By dynamically adjusting stimulation parameters (e.g., target location, intensity, frequency) based on real-time neural state estimates, the system could optimize transitions toward desired cognitive or affective states. This vision aligns with recent efforts to move beyond static stimulation protocols and toward adaptive, personalized, and feedback-driven neuromodulation strategies (Santaniello et al., 2011; Steffen et al., 2024). While we focused here on transitions between resting and sad mood states, the same framework can be applied to other cognitive or clinical states, and extended to new objective functions.

Finally, our methodology aligns with deterministic model-based RL approaches using latent dynamical systems (Hafner et al., 2025).

### Limitations

Despite the robustness of the modeling framework and the mechanistic insights it yields, the control approach remains to be validated. Specifically, we did not observe a significant association between control energy and mood ratings obtained during scanning, leaving open the question of whether the model-derived dynamics reflect subjectively experienced affective states. Furthermore, our inferences are based on full in silico simulations, which may not translate directly to practical neuromodulation: real-world stimulation protocols could impose additional physiological and technological constraints, and thus would likely require the inference of new, protocol-specific control strategies rather than a direct mapping of the simulated control energies. Future work should address these gaps by relating control metrics to behavioral or clinical measures and apply the proposed approach to infer stimulation strategies that account for the constraints of actual neuromodulatory interventions.

## 4 Conclusion

In summary, our study bridges the gap between dynamical systems theory, clinical neuroscience, and RL-based control theory by providing a data-driven framework to investigate neural state transitions and how they could be manipulated. The altered controllability and connectivity patterns in rMDD highlight the potential of this approach to uncover novel neural dynamics and therapeutic targets for mood disorders.

## 5 Methods

### 5.1 Sample

We reanalyzed a data set first reported in Zamoscik et al. (2014) (see also Huffziger et al., 2013; Lydon-Staley et al., 2019; Timm et al., 2017; Zamoscik et al., 2018; Zhang et al., 2023 for elaborate details). In brief, the study included 30 individuals with rMDD — defined as having experienced at least two episodes of MDD — and 30 HC matched for age, sex, and education, with no history or current diagnosis of MDD. The rMDD group had to be in partial or full remission for a minimum of two months. Exclusion criteria for all participants included clinical diagnoses of bipolar disorder, psychotic disorders, substance dependence, current substance abuse, generalized anxiety disorder, obsessive-compulsive disorder, post-traumatic stress disorder, eating disorders (all as per DSM-IV), and contraindications for MRI. A trained clinical psychologist conducted the diagnostic assessments using the Structured Clinical Interview for DSM-IV Axis I. The study received approval from the local ethics committee of Heidelberg University and conformed to the Declaration of Helsinki. All participants provided written informed consent.

### 5.2 Experimental Data

Based on the study by Zamoscik et al. (2014), we selected two 4.5-minute fMRI sessions for analysis: the first resting state and sad mood induction session from their more complex experiment. Sad mood induction was achieved using three personally significant negative life events, identified immediately prior to the fMRI session (e.g., a breakup, the death of a loved one, or a traumatic experience). These events were presented sequentially as keywords (each displayed for 1.5 minutes) accompanied by instrumental background music (excerpts from Albinoni’s Adagio in G minor). Mood assessments were conducted using the Positive and Negative Affect Scale (PANAS; Watson et al., 1988) before and after the sad mood induction. Both groups reported sadder mood after mood induction where rMDD reported a significantly more increased negative affect and a similarly decreased positive affect compared with HC (Zamoscik et al., 2014). During the resting state session, participants also viewed emotionally neutral words (namely, “rest 1”, “rest 2”, “rest 3”).

#### 5.2.1 Preprocessing

The study used a 3T Trio Tim Scanner with a 12-channel head coil (Siemens Healthineers, Erlangen, Germany), acquiring functional images at a rate of 1.5s (TR). For additional details on data acquisition, data processing, and artifact correction, see Section A.1.

To obtain time-series data for modeling, anatomical masks were used to extract 11 brain regions from both hemispheres (Figure 2a). These regions were chosen to provide system-level coverage of five large-scale networks consistently implicated in major depressive disorder—salience/limbic, default mode, cognitive control, reward, and memory networks—based on converging evidence from metaanalytic studies showing structural and functional alterations in these circuits Gou et al., 2023; Kaiser et al., 2015; Ng et al., 2019; Schmaal et al., 2016.

Time series were standardized and subsequently bandpass filtered using cutoff frequencies of 0.0083 Hz and 0.15 Hz. For each subject, data was projected onto the top three PCs, which explained an average of 76% of the variance, with a minimum of 50% across all subjects. This yielded a 66-dimensional time series (2 hemispheres x 11 regions x 3 PCs), comprising 320 time points per subject, with 160 time points for each of the resting state and mood induction conditions. Finally, the time series — denoted as *x*_*t*_ in the following — were standardized for PLRNN, SVM, and GMM training. This scaling was reversed during control optimization, restoring each dimension to its original variance.

### 5.3 DSR Model and Control

#### 5.3.1 DSR Model Architecture

To model the dynamics within the selected ROIs, we employ a state-space framework comprising a latent dynamical model coupled with an observation model. The latent dynamics are governed by the shallow PLRNN (shPLRNN; Hess et al., 2023), defined as

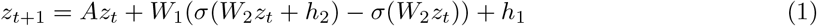

with latent states *z*_*t*_ ∈ ℝ^*M*^, diagonal matrix *A* ∈ ℝ^*M* ×*M*^, connectivity matrices *W*_2_ ∈ ℝ ^*L×M*^ and *h*_2_ ∈ ℝ^*L*^, and biases *h*_1_ ∈ ℝ^*M*^, where *L* is the dimension of the hidden layer, and *σ*(·) is an elementwise piecewise linear activation function (the Rectified Linear Unit; ReLU).

The observation model is linear and defined as 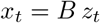, where *B* ∈ ℝ^*O*×*M*^ and *O* = 66 is the observation dimension.

Each model is trained jointly on both the resting state and sad mood induction data from a single subject, without access to condition labels during training.

#### 5.3.2 Model Training

Models were trained using backpropagation through time (BPTT) combined with generalized teacher forcing (GTF; Hess et al., 2023), a technique in which model-generated latent states are partially realigned with encoded states after gradient computation using linear interpolation with the interpolation parameter decaying from *α*_0_ to *α*_1_ during training. This alignment helps mitigate exploding and vanishing gradients during training. Encoded latent states *z*^enc^ were obtained by projecting observations into the latent space via the Moore-Penrose pseudoinverse *B*^+^.

Training was performed by minimizing a regularized mean squared error loss, which penalizes discrepancies between model-generated and empirical observations, as well as between model-generated and encoded latent states,

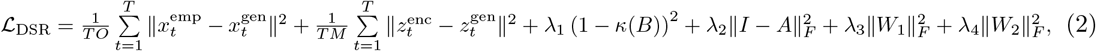

where 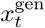 and 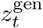 refer to model generated observations and latent states generated according to eqn. (1) and the observation model, and *κ*(*B*) denotes the condition number of the observation matrix *B* that is regularized towards 1 to promote numerical stability of the pseudoinverse via *λ*_1_. The *λ*_2_-term regularizes the diagonal entries of *A* toward 1 and *λ*_3_ and *λ*_4_ regulate *L*_2_ penalties on *W*_1_ and *W*_2_, ∥ · ∥_*F*_ denotes the Frobenius norm.

We use the RAdam optimizer (Liu et al., 2020), with an exponentially decaying learning rate from *η*_0_ to *η*_1_, and momentum parameters *β*_1_ = 0.9 and *β*_2_ = 0.999.

The models are trained on the first 120 time steps of each session (75%; training set) and evaluated on the remaining 40 time steps (test set). During each training epoch, a full batch of 162 sequences of length *T* = 40 is sampled—comprising 81 sequences from the 120 training time steps of both the resting state and sad mood induction sessions. The hyperparameters including the matrix dimensions used in training were selected via grid search on the test set for a subset of five subjects, with the final values summarized in Table 1. For each participant, we trained five models with Xavier initialization (Glorot and Bengio, 2010) for up to 200,000 epochs and selected the model with the lowest test loss for subsequent experiments.

**Table 1:** Hyperparameters used for DSR model training.

| $M$ | $L$ | $\alpha_0$ | $\alpha_1$ | $\beta_1$ | $\beta_2$ | $\lambda_1$ | $\lambda_2$ | $\lambda_3$ | $\lambda_4$ | $\eta_0$ | $\eta_1$ |
| --- | --- | --- | --- | --- | --- | --- | --- | --- | --- | --- | --- |
| 66 | 15 | $10^{-1}$ | $10^{-3}$ | 0.9 | 0.999 | $10^{-1}$ | 0 | $10^{-4}$ | $10^{-4}$ | $10^{-3}$ | $10^{-4}$ |

##### Physiological Confounds

Since elevated connectivity levels were observed in the rMDD group—and such effects can potentially arise from physiological artifacts (e.g., cardiac and respiratory signals, Wilding et al., 2024; Zamoscik et al., 2018) — we re-estimated all shPLRNN models after removing these artifacts from the data. Using the TAPAS PhysIO toolbox (Kasper et al., 2017), we extracted subject-specific cardiac and respiratory regressors from recorded physiology time series. We then applied ridge regression (*l*_2_ regularization) to model and subtract these nuisance signals, optimizing the regularization weight via cross-validation and assessing fit on a held-out test set. Importantly, the elevated connectivity patterns not only persisted but in some instances became even more pronounced (Figure A9). To remain conservative and retain the original neural signal structure, all subsequent analyses therefore used the models trained on the uncorrected data.

#### 5.3.3 DSR Model Evaluation

To quantify how well the DSR model reproduces the temporal structure of the data, we compare the power spectrum of each latent dimension between the measured trajectory and the model’s reconstruction along the entire time series. Concretely, for each dimension *i* ∈ { 1, …, *O* } we compute the Fast Fourier transform of the time series (Cooley and Tukey, 1965), smooth the resulting power spectrum with a Gaussian kernel, and renormalize it to obtain two spectral densities, *S*_*i*_(*ω*) (measured) and *P*_*i*_(*ω*) (reconstructed). We then summarize their discrepancy by the mean Hellinger distance, 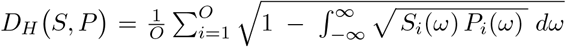. This metric ranges from 0 (identical spectra) to 1 (maximally dissimilar), and thus provides a dimension-averaged measure of how faithfully the shPLRNN captures the frequency content of the original signals.

To further assess model performance on unseen data, we also compared the functional connectivity patterns in the empirical and reconstructed test-set trajectories. Specifically, we computed the Pearson correlation matrix across all observed regions for both the empirical data and the held-out shPLRNN reconstructions. This connectivity comparison ensures that the shPLRNNs not only reproduce spectral properties but also preserve the inter-regional coupling structure present in the test data.

#### 5.3.4 Model Predictive Control

Our MPC framework is an RL strategy that leverages a model of the system to predict future dynamics and optimize control actions over a finite time horizon. Instead of estimating a value function that describes the expected return from a state, our framework implicitly evaluates states by running forward simulations. At each time step, it solves an optimization problem to minimize a cost functional — applying only the first control step before re-optimizing — and thereby enabling real-time adaptation. We integrate MPC with DSR to determine the stimulus required to steer the brain into desired states (Figure 1). Using shPLRNNs (eqn. 1) trained on empirical fMRI data, we define an additive, state-dependent control term *u*(*z*_*t*_) that modifies the latent dynamics as follows

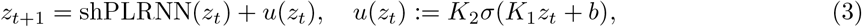

where σ(·) = ReLU, *K*_1_ ∈ ℝ ^*Q×M*^, *K*_2_ ∈ ℝ^*M×Q*^ and *b* ∈ ℝ^*Q*^. The shPLRNN parameters are fixed post-training; only the control parameters (*K*_1_, *K*_2_, *b*) are optimized (Figure A1).

##### Soft Target Objective

To allow flexible control, we define a soft target using a GMM trained on empirical resting state or sad mood induction data (see Section 5.3.6). Instead of reaching a fixed state, the controlled trajectory is guided into high-likelihood regions under the GMM.

The loss function thus consists of a negative log-likelihood term and a regularization term penalizing control energy:

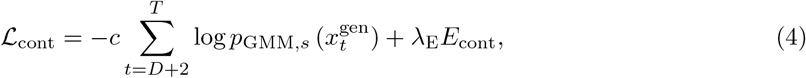

where 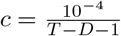 ensures loss terms are comparable in magnitude, and control energy is defined as

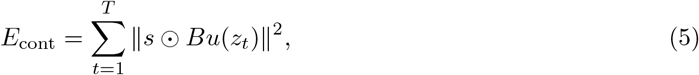

where *s* are the pre-standardization standard deviations, restoring the original scale of each PC.

##### Control Optimization

Control parameters *K*_1_, *K*_2_, *b* are optimized using the RAdam optimizer over up to 30,000 epochs with a constant learning rate *η*. Each trajectory starts from an empirical state and runs for *T* = *D* + 11 steps (1 for initialization, *D* before loss evaluation, and 10 to maintain the target). We use 240 empirical states for training and 80 for testing. Hyperparameters were set to *λ*_E_ = 0.1, *Q* = 17, and *η* = 0.01. Among five independently trained models, the one with lowest test loss is selected for analysis. The optimization was robust to initialization (Figure A10).

##### Restricting Control to Specific Brain Regions

To assess controllability and region-specific influence, we constrain the control to act only on selected brain regions. While latent space constraints are trivial (e.g., via L1-regularization to promote sparsity), enforcing constraints in observation space is more complex due to the linear observation model, where latent dimensions are mixed and do not map directly onto individual brain regions.

We overcome this by projecting *u*(*z*_*t*_) onto the null space of the decoder rows corresponding to regions where control should be inactive. Let 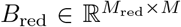 denote the submatrix of *B* corresponding to these regions. We compute an orthonormal basis 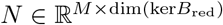 of ker(*B*_red_) via singular value decomposition and construct the projection matrix *P* = *NN* ^T^ (see also Fechtelpeter et al., 2025). The constrained dynamics then become

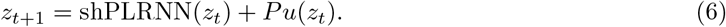

This ensures that *Bu*(*z*_*t*_) has no component in the restricted observation dimensions. The projection matrix *P* is computed once before training and remains fixed throughout optimization.

#### 5.3.5 Coupling Matrix and Lyapunov Spectra

The shPLRNNs (eqn. (1)) permit efficient computation of coupling matrices and Lyapunov spectra along individual trajectories by exploiting the model’s Jacobians 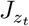 . At latent state *z*_*t*_, the Jacobian of the update map *z*_*t*_ ↦ *z*_*t*+1_ is

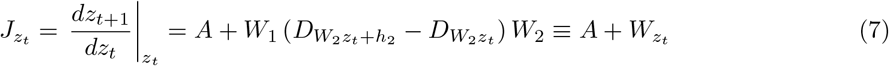

where

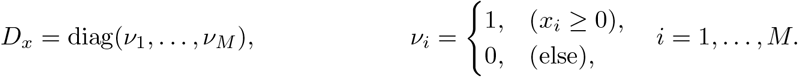

Here 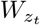 captures how different latent dimensions interact at time *t*, and is often called the cross-connection matrix.

Since our observations *x*_*t*_ are related to *z*_*t*_ by the linear map 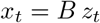, the corresponding Jacobian in observation space is

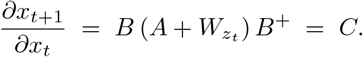

In other words, *C* is the coupling matrix of the observable dynamics. We thus interpret ∥*C*∥(its spectral norm) as quantifying how strongly activity in one region influences activity in another. We also evaluate ∥*C*∥ at every _*zt*_ visited by the shPLRNN when driven by empirical resting state and sad mood-induction data, yielding a time-resolved measure of inter-regional coupling.

To compute Lyapunov exponents, we apply the algorithm of Vogt et al. (2022), which orthogonally projects 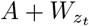 onto tangent subspaces along each trajectory.

#### 5.3.6 Statistical Analysis

##### State Separation

To estimate the empirical distribution of brain states during resting state and sad mood induction, we trained separate GMMs of the respective datasets per subject. GMMs provide a probabilistic description of the state-space, capturing regions of high empirical density while allowing generalization to nearby states. Model selection based on the Bayesian Information Criterion (BIC) and Akaike Information Criterion (AIC) indicated that six mixture components best described the data (Figure A2b).

To quantify group-level differences in brain state distributions, we computed three distances between resting and sad mood GMMs: (i) maximum mean discrepancy (MMD), (ii) the *L*^2^ norm in the space of square-integrable functions (both computed in closed form), and (iii) the sliced Wasserstein distance (estimated via sampling).

##### Control Evaluation and Group Statistics

For model evaluation under control, trajectories were initialized from each of the 160 empirical brain states (resting or sad mood induction). After an initial warm-up period of *D* steps, we computed both the NLL under the target GMM and the cumulative control energy.

To assess regional specificity, the control experiment was repeated 11 times — each time with control restricted to one bilateral region. Across all 30 subjects per group and 160 initializations per condition, the resulting trajectories span *T* = 80 time steps, yielding NLL and energy tensors of shape (30, 160, *T*). For statistical analysis, the first *D* steps were excluded to limit analyses to time points where target distance was penalized during training; the median was then computed across time and trials to yield a single value per subject.

Outliers were excluded based on the 1.5×interquartile range criterion. Group differences were assessed using the Mann-Whitney U test (two-sided). All *p*-values were corrected for multiple comparisons using the Benjamini-Hochberg procedure to control the false discovery rate (FDR).

## Acknowledgments

This work was funded within Germany’s Excellence Strategy EXC 2181/1 – 390900948 (STRUCTURES), the German research center TRR 265, subproject A06, and the Hector II foundation.

## A Supplementary Methods

### A.1 fMRI Preprocessing

#### Imaging Parameters

Functional imaging was conducted with T2*-weighted EPI scans (TR = 1.5 s, flip angle *α* = 80°, TE = 28 ms, 24 slices, voxel size 3 × 3 × 4 mm3, 180 volumes per phase), while high-resolution anatomical images were captured using T1-weighted MPRAGE sequences (TR = 2.3 s, TE = 3.03 ms, voxel size 1 × 1 × 1 mm^3^). Physiological data, namely pulse and respiration, were recorded at 50 Hz. The first 20 of the 180 images per phase were discarded and preprocessing was conducted with SPM12 (v7738).

#### Data Preprocessing and Artifact Correction

Preprocessing for the anatomical images included segmentation of the anatomical images and normalization to the ICBM 2009b Nonlinear Asymmetric template, and for the functional images slice timing correction, realignment, coregistration to the anatomical data, normalization to standard space, and spatial smoothing with a FWHM=8x8x8 mm^3^ Gaussian kernel.

Onset effects for the key words were modeled by convolving their onset stick function with the canonical hemodynamic response function and included in a regression model further containing the six standard motion parameters, their derivatives, the squared values of them, white matter, cerebrospinal fluid, and global signals (Parkes et al., 2018) as well as dummy regressors for volumes affected by small movements (framewise displacement *>* 0.5mm, global intensity change *z >* 4). We removed these confounds via ridge regression, then extracted the residual time series from each target region and applied the preprocessing pipeline described in Section 5.2.1.

### A.2 DSR Hyperparameters

## B Supplementary Figures

**Figure A1:**
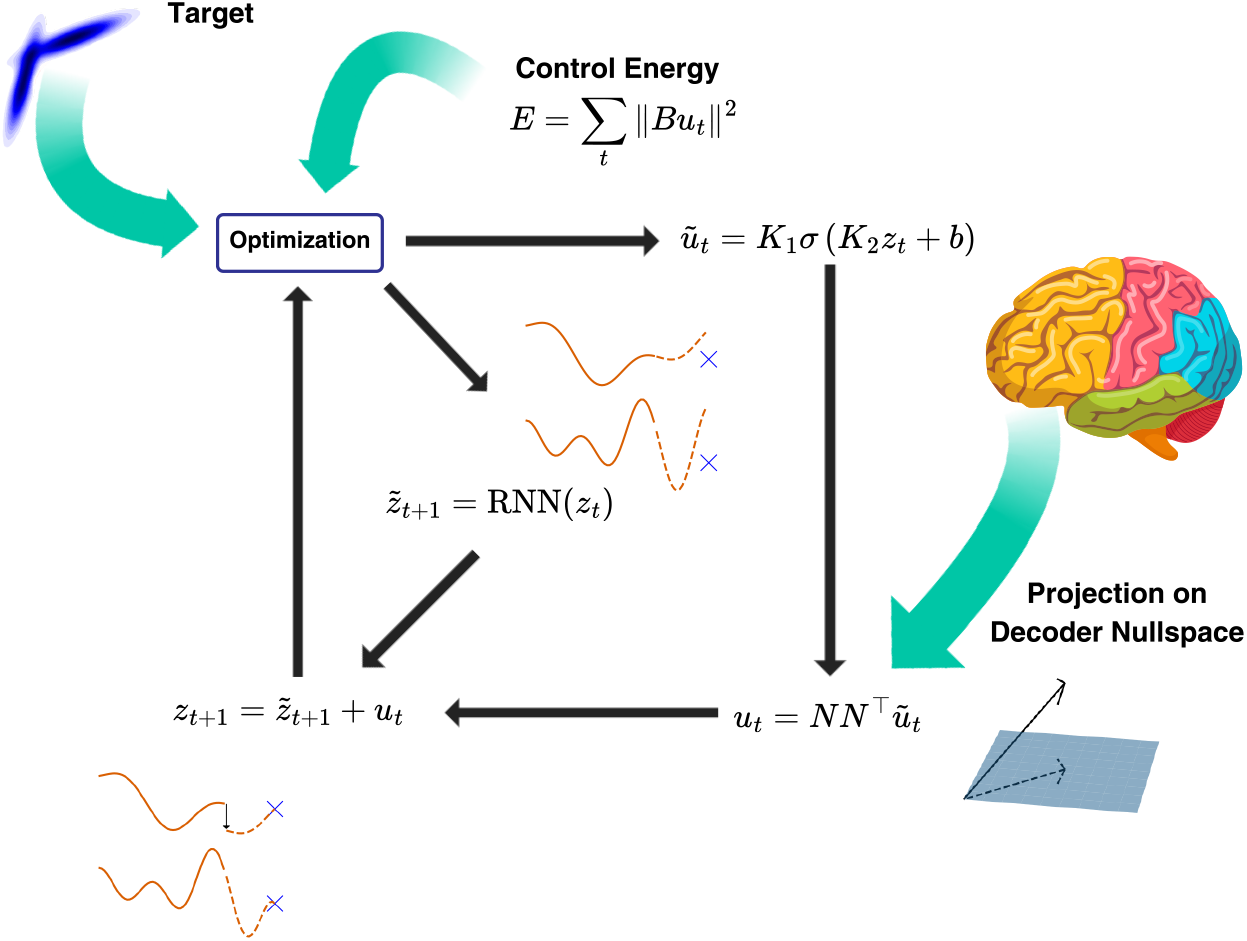
Closed-loop MPC Framework. shPLRNNs inferred on the data model the uncontrolled brain dynamics (center). Control is applied via an additive term *u*_*t*_ = *u*(*z*_*t*_) (bottom left), optimized to drive the system toward a target state while minimizing control effort (top left). To restrict control to specific brain regions, *ũ*_*t*_ is projected onto the null space of corresponding rows of the decoder *B*, using the projection matrix *NN* ^T^ (bottom right).

**Figure A2:**
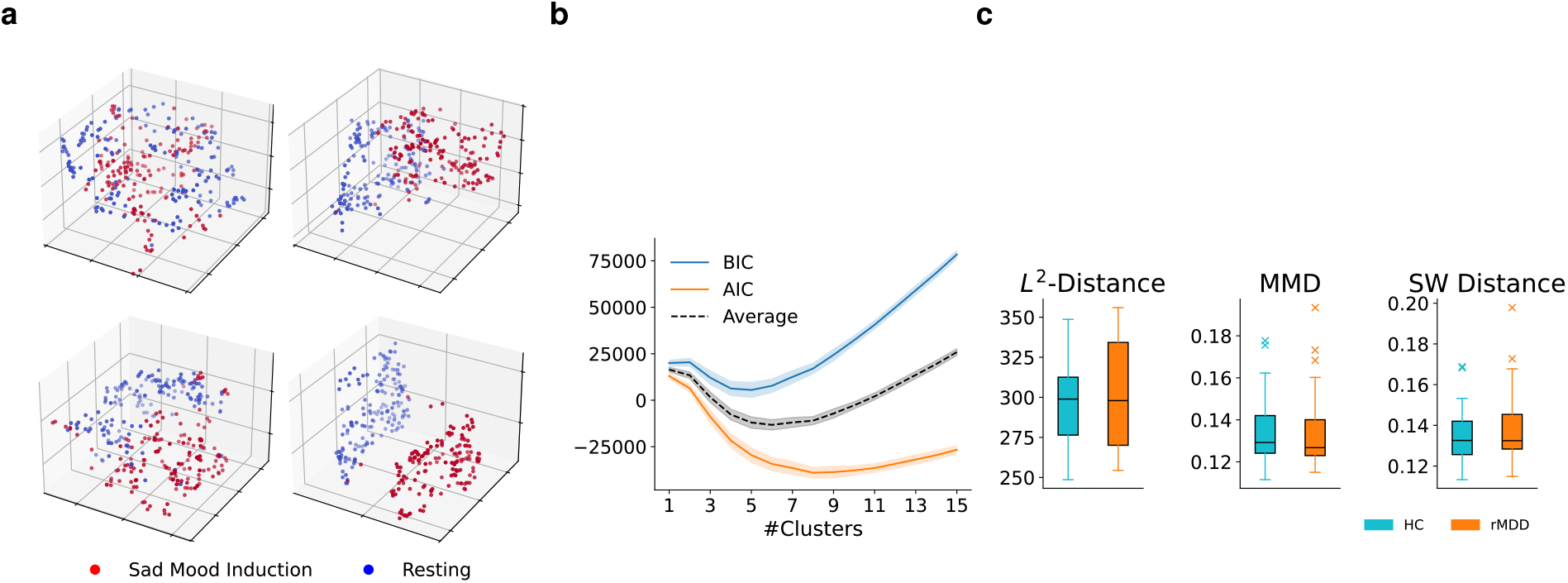
**a:** UMAP dimensionality reduction applied to states sampled from the Gaussian Mixture Models (GMMs) confirm that the empirical separation is well preserved (cf. Figure 2b). **b:** Model selection using AIC and BIC across subjects suggests optimal cluster number of 6. **c:** Group comparisons of the distance between GMMs fitted to the resting and sad mood induction states. No significant differences were observed across metrics: *L*^2^ Distance (*p* = 0.81), Maximum Mean Discrepancy (MMD, *p* = 0.92), and Sliced Wasserstein Distance (SW, *p* = 0.74).

**Figure A3:**
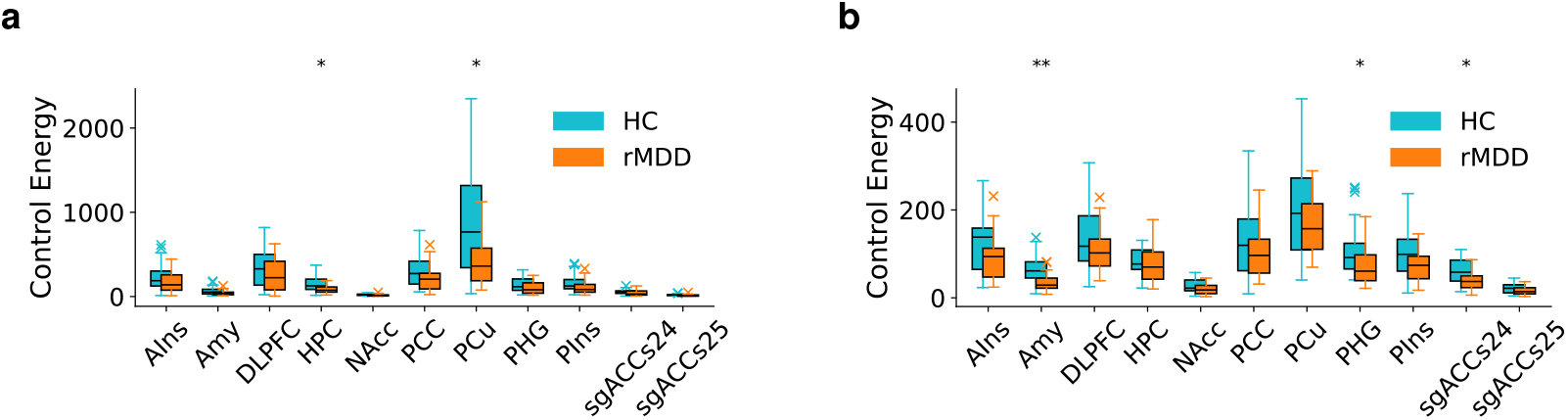
Robustness to Variations in Control Energy Regularization *λ*_E_ and Time Horizon *D* for Control toward Sad Mood. **a**: Without an explicit penalty on control energy (*λ*_E_ = 0), overall energy levels rise, and the rMDD group generally requires less control (statistically significant within HPC and precuneus; PCu). **b**: With a shorter horizon (*D* = 5), the model has fewer time steps before penalties on control energy and target distance apply; rMDD likewise requires less control energy.**a, b** ^*^*p*_FDR_ *<* 0.05.

**Figure A4:**
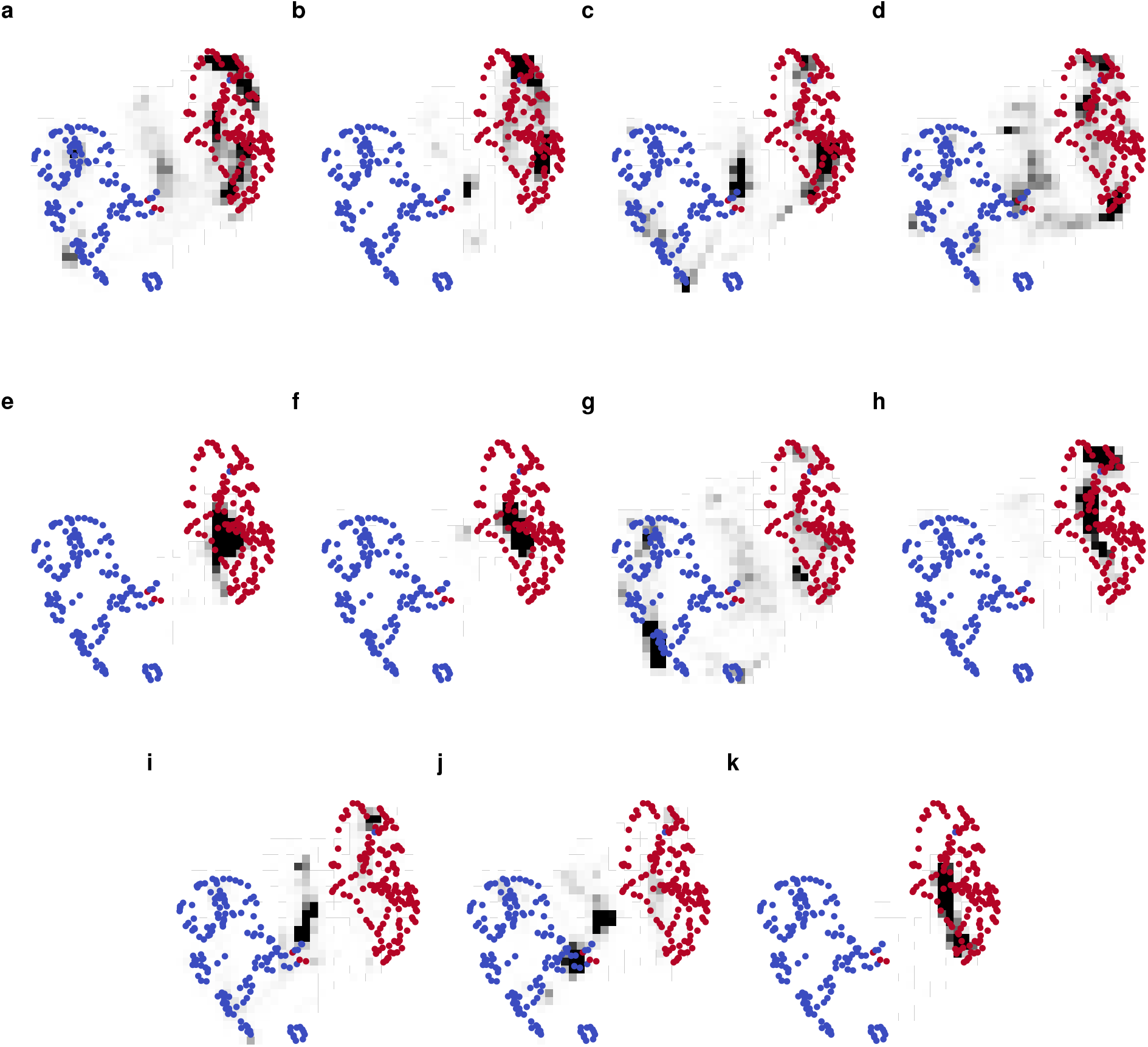
**a-k**: Two-dimensional UMAP visualizations of empirical states (sad mood induction in red, resting in blue) and trajectories (black) controlled torwards sad mood for different targeted brain regions in a fixed subject. The control successfully maintains activity within the sad mood induction state-space region. Trajectories from multiple initializations are aggregated by spatial binning, and relative frequencies per bin are displayed as a heatmap. Time steps smaller than *D* = 10 were discarded.

**Figure A5:**
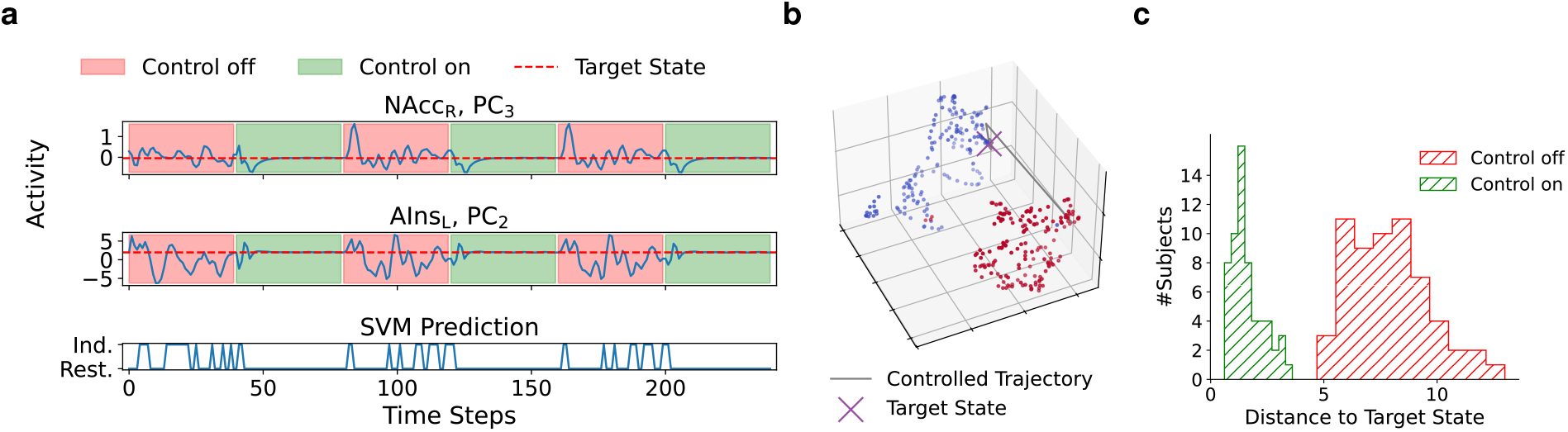
Hard Target Control: A specific sad mood state (red dashed line) was designated as the target, and control was optimized using mean squared error (MSE) loss. Control was applied to all cortical regions. **a**: When control is activated, trajectories reliably converge toward the target state (red dashed line); when deactivated, dynamics revert to chaotic behavior. This effect is visible in both controlled (right AIns) and uncontrolled (left NAcc) dimensions. The SVM decoder confirms convergence into the resting state region. **b**: UMAP visualization of a trajectory initialized at sad mood induction (red) converging to the target resting state (blue). **c:** Control significantly reduces the Euclidean distance to the target state (median across time), consistently across all subjects.

**Figure A6:**
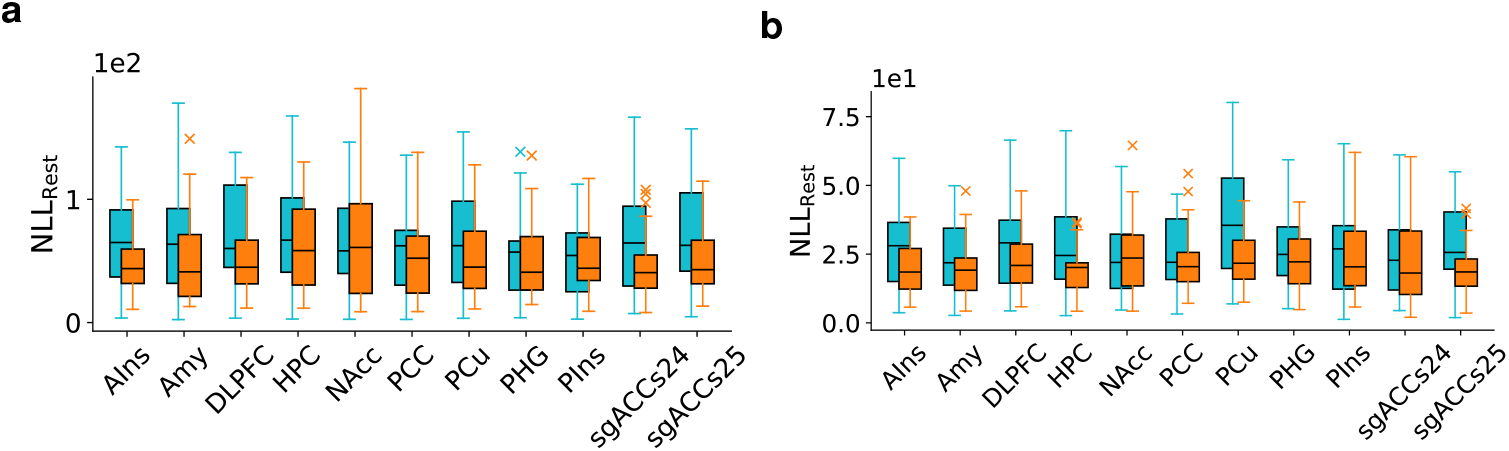
No Group Differences in Terms of Proximity to Resting States: Both for the transition from *resting state to sad mood* (**a**) and *sad mood to resting state* (**b**), rMDD and HC maintain a similar proximity to the resting state.

**Figure A7:**
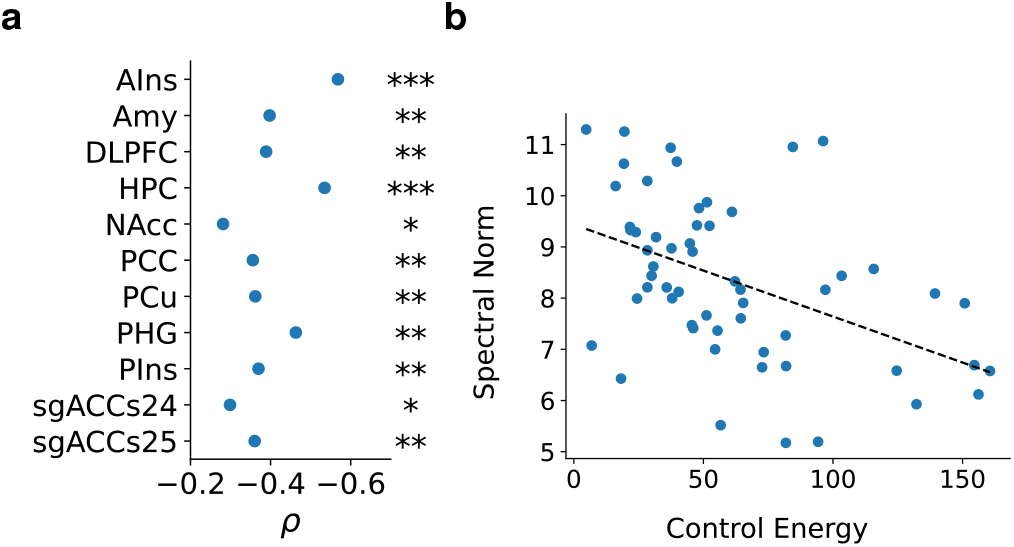
Correlation Transitions Toward Resting State. **a**: For the control in the resting direction, the negative correlation between region coupling and control energy again shows that increased coupling strength is associated with regions with reduced energy expenditure. **b**: Specific values of the spectral norm and the energy for each subject on the example of the PHG (c.f. Figure 5c and 5d).

**Figure A8:**
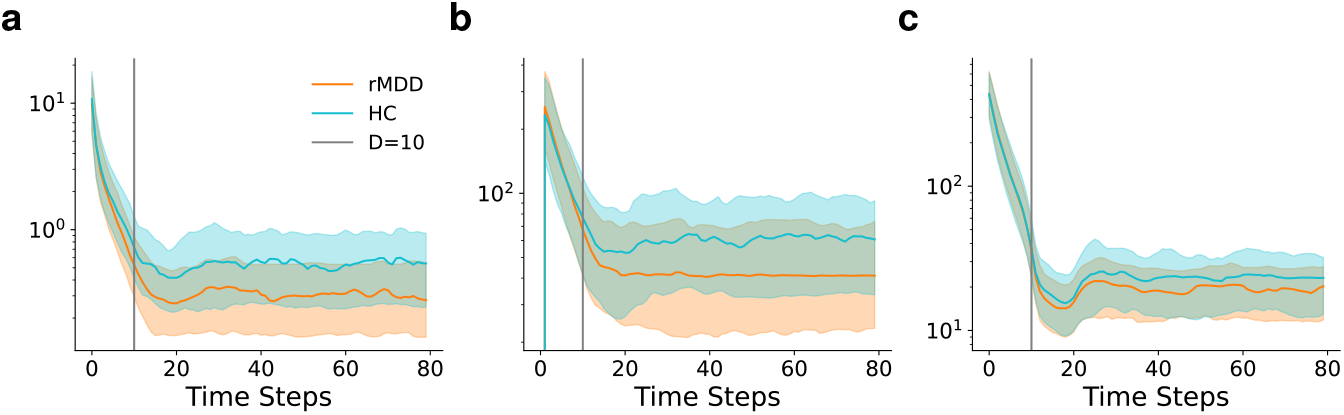
Temporal Evolution of the Evaluation Measures. Control energy (**a**), NLL_Rest_ (**b**), and NLL_Ind_ (**c**) for the case of amygdala-targeted control. Curves show the median across subjects and initializations; shaded areas denote the inter-quartile range. While rMDD subjects consistently require less control energy, no significant differences are observed in the likelihoods that quantify the proximity to initial and target states.

**Figure A9:**
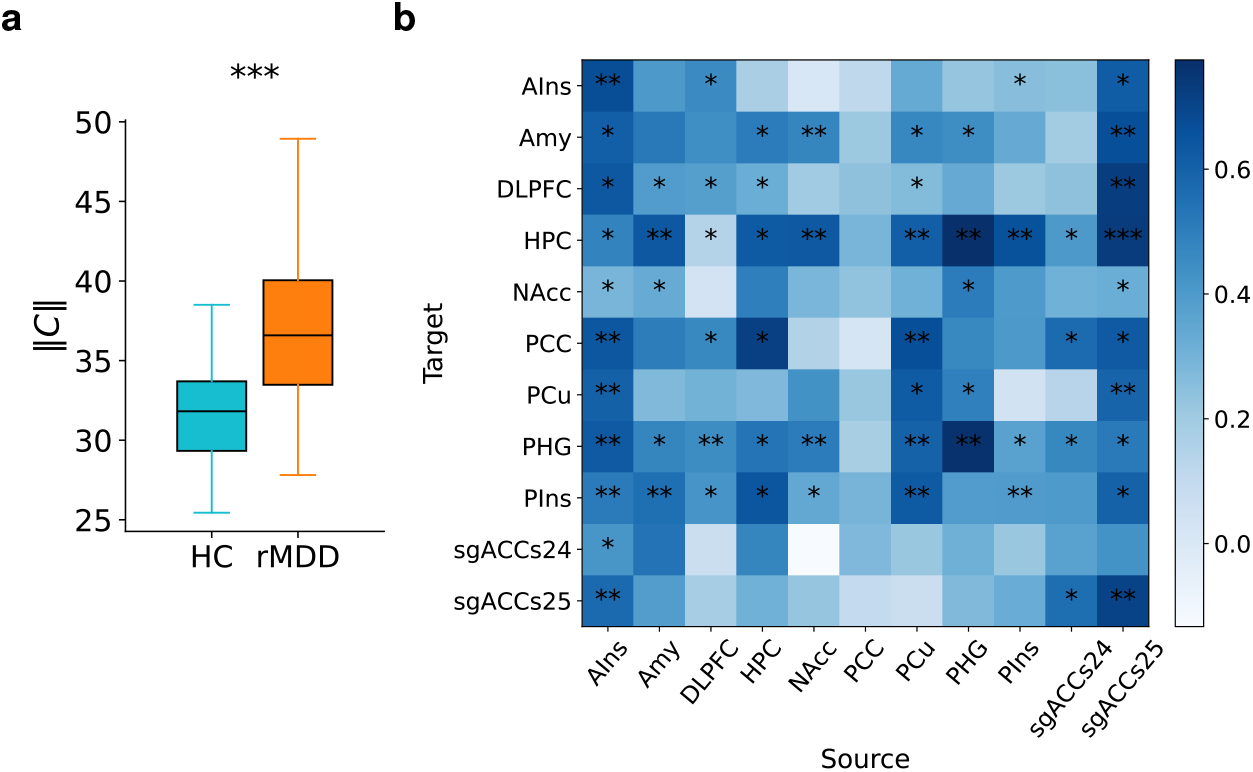
Connectivity Differences When Training was Performed on Data Corrected for Potential Physiological Artifacts. When correcting the data for physiological effects (i.e. cardiac and respiratory noise) before training the shPLRNNs, the significant difference in connectivity between the groups remains evident. Both the global coupling (**a**) and the pairwise connectivity between brain regions (**b**, difference in median, rMDD-HC) is significantly stronger in rMDD. Here, ^*^*p*_FDR_ *<* 0.05,^**^ *p*_FDR_ *<* 0.01,^***^ *p*_FDR_ *<* 0.001.

**Figure A10:**
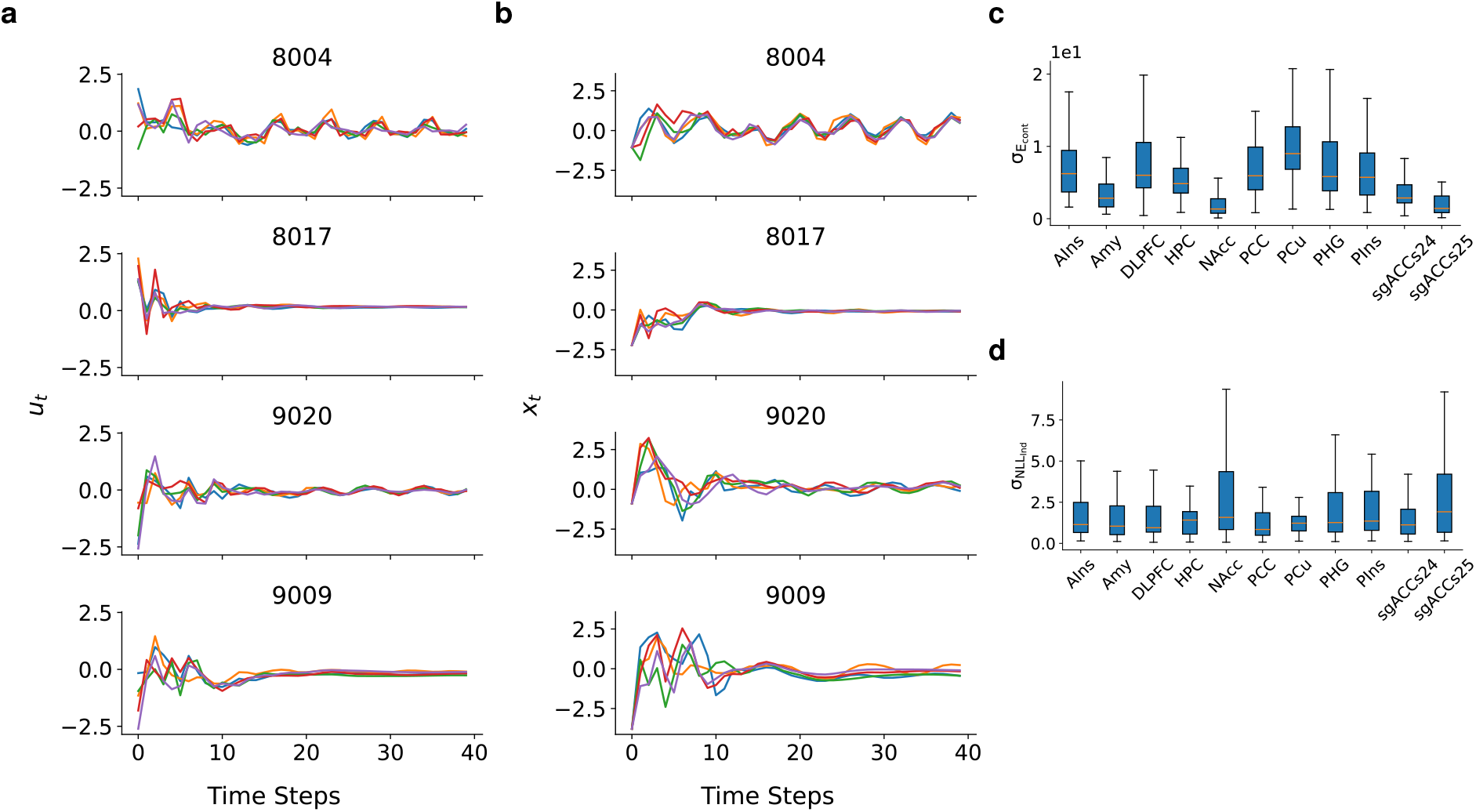
Robustness of Control Parameters to Initialization. For each experiment and subject, five models were independently optimized. Controlled trajectories show predominantly consistent control inputs *u*_*t*_ (**a**) and corresponding activity *x*_*t*_ (**b**) across runs, each represented by a separate color-coded curve, displayed for a selected number of subjects. While the chaotic dynamics amplify small deviations over time, variance across runs of control energy (**c**) and NLL_Ind_ (d) remains substantially smaller than inter-subject variance (cf. Figure 3e, 3f). The models with the lowest loss were used for analyses.

